# A machine learning method for estimating the probability of presence using presence-background data

**DOI:** 10.1101/2022.03.29.486220

**Authors:** Yan Wang, Chathuri L. Samarasekara, Lewi Stone

**Author notes:** The first two authors contributed equally to this paper.

## Abstract

Estimating the prevalence or the absolute probability of presence of a species from presence-background data has become a controversial topic in species distribution modelling. In this paper we propose a new method by combining both statistics and machine learning algorithms that helps overcome some of the known existing problems. We have also revisited the popular but highly controversial Lele and Keim (LK) method by evaluating its performance and assessing the RSPF condition it relies on. Simulations show that the LK method with unfounded model assumptions would render fragile estimation/prediction of the desired probabilities. Rather we propose the local knowledge condition, which relaxes the pre-determined population prevalence condition that has so often been used in much of the existing literature. Simulations demonstrate the performance of the CLK method utilising the local knowledge assumption to successfully estimate the probability of presence. The local knowledge extends the local certainty or the prototypical presence location assumption, and has significant implications for demonstrating the necessary condition for identifying absolute (rather than relative) probability of presence without absence data in species distribution modelling.

## 1 Introduction

Ecologists and environmental scientists make use of species distribution models (SDMs) to help understand how organisms selectively use their resources and the factors which shape their spatial distributions. SDMs have therefore become fundamental for many tasks in modern ecological modelling (Elith et al., 2006; Lele and Keim, 2006; Phillips and Elith, 2013). Considerable research has been carried out on species distribution modelling, as outlined in the extensive literature review of Guillera-Arroita et al. (2015). Of the different types of spatial data used in SDMs, Presence Background (PB) data are plentifully available and easy to access. PB data consists of a list of “presences” or locations where individuals/species have been observed, but there is no information about locations of absences (Guillera-Arroita et al., 2015). This latter characteristic makes the data difficult to work with. PB data are often available from museum, herbarium collections and other historical database records. It now becomes increasingly available via citizen science projects and online repositories such as the Global Biodiversity Information Facility (GBIF: http://www.bgif.org). This paper will focus on the methodologies developed that relate to PB data, which has been used in approximately 50% of SDM papers, according to the survey in Guillera-Arroita et al. (2015). When using PB data, the key objective is usually to accurately estimate the site-specific probability of presence in a geographical area.

Numerous methods have been developed for modelling PB data with SDMs, including statistical regression models known in the literature as the LI (Lancaster and Imbens, 1996), LK (Lele and Keim, 2006), Expectation-Maximization (EM) (Ward et al., 2009), Scaled Binomial Loss Model (SB) (Phillips and Elith, 2011), spatial point process models (PPMs) (Warton and Shepherd, 2010; Renner and Warton, 2013), machine learning (ML) methods such as MAXENT (Phillips et al., 2006), presence and background learning (PBL) algorithm (Li et al., 2011) and boosted regression trees (Elith et al., 2008), and various other methods. It is commonly recognized that the actual (or absolute) probability of presence given environmental covariates, namely the resource selection probability function (RSPF), cannot be predicted from presence-only or PB data, when there is no extra information or conditions available to make use of (Elith et al., 2008; Hastie and Fithian, 2013; Phillips and Elith, 2013; Wang and Stone, 2019; Ward et al., 2009). Without the extra information, all of these methods have been shown only to be useful in their ability to estimate the ratio between the probability of presence and the probability of prevalence, also known as the resource selection function (RSF) (Manly et al., 2002) or the “relative” probability of presence (Wang and Stone, 2019). In fact, the conditions required to estimate the true conditional probability of presence from PB data has become a highly controversial topic (Lele and Keim, 2006; Hastie and Fithian, 2013; Knape and Korner-Nievergelt, 2015; Phillips and Elith, 2013; Solymos and Lele, 2015; Wang and Stone, 2019; Ward et al., 2009). LK claimed their method can successfully achieve this goal when the so called ‘RSPF conditions’ are satisfied (Lele and Keim, 2006). (We will outline these conditions shortly.) A paper devoted to this question (Solymos and Lele, 2015) argues that the class of admissible models that satisfy RSPF conditions is very broad and does not excessively restrict the application of the LK method. However, this contradicts arguments in many other papers including the important paper of Hastie and Fithian (2013).

Wang and Stone (2019) recently revealed the close connection between many commonly used but seemingly disparate methods, such as the LI, LK, EM, SB, MAXENT and PPMs. In particular, Wang and Stone (2019) proposed a new unified Constrained LK (CLK) method, which serves as a generalisation of the better known existing approach. However, the CLK method requires information of the population prevalence as the input, which is difficult to obtain or estimate in practice. This renders the CLK method, along with other methods (such as the SB and EM methods) that also require prior information of the population prevalence, limited in their practical applications.

In this paper, we propose a “refined” CLK method that can be used to accurately estimate the population prevalence so that the probability of presence can be ultimately estimated. A key assumption of the method is that there exists ‘local knowledge’ where habitats are maximally or partially suitable for a species. This means there must be some sites where we have knowledge about the resource selection probability of a species. The idea was inspired by the work of Elkan and Noto (2008) and Li et al. (2011). Elkan and Noto (2008) originally proposed the presence background learning (PBL) algorithm for the positive and unlabelled problem in the field of machine learning. Li et al. (2011) further developed the algorithm to estimate the actual probability of presence in modelling species’ distributions.

The performance of the method proposed here has been tested and compared with the popular LK method through extensive simulation studies, where a wide range of resource selection probability functions were taken directly from Solymos and Lele (2015) to ensure that the RSPF conditions were satisfied according to their definitions in Solymos and Lele (2015).

Our simulation studies reveal that the performance of the LK method is often poor even in situations where the RSPF condition is satisfied, in contrast to the problematical claims of Solymos and Lele (2015). On the other hand, simulations demonstrate the excellent performance of the proposed method when what we term ‘local knowledge’ is satisfied. The experiments show that the ”local knowledge” condition is not just helpful but is necessary for identifying probability of presence from presence-background data. Compared to the commonly used pre-determined population prevalence (Ward et al., 2009; Phillips and Elith, 2011; Wang and Stone, 2019), the “local knowledge” relaxes the required information and is thus less constrained.

## 2 Materials and Methods

### 2.1 Description

When working with SDMs, we are interested in whether a species is present or absent at a particular site, conditional on environmental covariates (denoted by *x*). Here the variable *y* = 1 represents the species’ presence while *y* = 0 represents its absence at a particular site. More specifically, a key goal is to estimate the conditional probability of Pr(*y* = 1|*x*), namely the absolute probability of presence at a site, based on the covariate *x* measured at that site. The overall population prevalence will be denoted as *π* = Pr(*y* = 1), i.e., fraction of sites in the study area in which the species is present. A common practice in statistical modelling is to assume a parametric structure for Pr(*y* = 1|*x*), for example, the logit function,

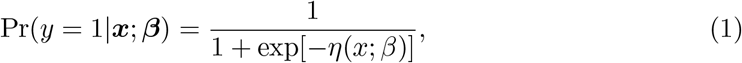

where *η*(*x*; *β*) can be a linear or nonlinear function of *x*, and *β* are parameters that need to be estimated.

The form of the parametric function is critically important for some statistical methods in SDM such as the LK method (Lele and Keim, 2006), and can determine the success of the method. In contrast, our proposed model is less reliant on the explicit form of the parametric function. Examples will be provided in section 2.6 to show the robustness of our proposed method.

The presence-background data can be viewed according to the following summary which is made use of throughout the paper. Let *P* be the set of presence samples which contains *p*_1_ observed presence data points (which in ML is referred to as “labelled data”). *B* is a set of *n*_0_ background samples in which each sample’s presence or absence is unknown (which in ML is referred to as “unlabelled data”). Assume the background samples contain *p*_2_ true presences although they have not been observed and are unknown. We use *s* to represent the sampling stratum, with *s* = 1 for the “labelled” samples appearing in *P* and *s* = 0 for “unlabelled” samples in *B*. If *s* = 1, it is known that *y* = 1 and that there is a “presence”. But, if *s* = 0 (i.e., in the background data), we do not know whether *y* = 1 or *y* = 0. Pr(*s* = 1) gives the probability of a species being observed or the probability of belonging to the presence samples (i.e. being labelled in the language of ML).

### 2.2 Recap of the LK and CLK methods

The LK method defines the target log likelihood of the presence samples as (Lele and Keim, 2006);

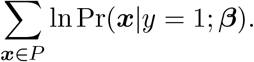

By applying Bayes rule it becomes;

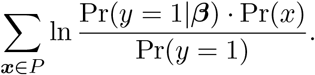

Dropping the terms that do not depend on the *β*, we obtain;

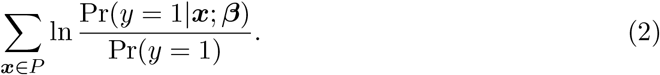

Lele and Keim (2006) used the background data *B* to estimate the denominator term Pr(*y* = 1) = *π* of population prevalence empirically, and obtained the following log-likelihood.

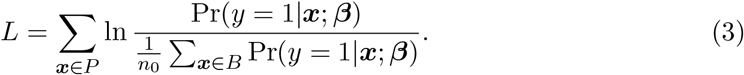

In Eqn. 3, the species’ prevalence has been approximated by the average probability over the background samples (Hastie and Fithian, 2013; Phillips and Elith, 2013; Wang and Stone, 2019). In Lele and Keim (2006), standard optimisation techniques are used to find the coefficients ***β*** that maximizes Eqn. 3.

Lele and Keim (2006) and Solymos and Lele (2015) argued that the LK method is valid in estimating the absolute probability of presence, if the so-called RSPF conditions hold. When stated simply, the RSPF conditions include 1) the true (actual) function of log Pr(*y* = 1|*x*) should be non-linear, and 2) not all covariates can be categorical. Readers are referred to Solymos and Lele (2015) for a complete discussion of these conditions. The logit function given in Eqn. 1 is one of the parametric functions that meet the RSPF conditions. (The complementary log-log (cloglog) function is another example that meets the conditions.) When these conditions are satisfied, it is claimed that the LK method can estimate the absolute probability of presence of a species without any other extra information needed (Lele and Keim, 2006; Solymos and Lele, 2015). Despite this, Hastie and Fithian (2013) have given an in depth analysis showing why the LK method is incapable of estimating the correct parameter values when data arise via models that are nearly linear on the logit scale.

Existing literature makes it clear that information of population prevalence *π* is required in advance to estimate the absolute probability of presence at each site (Elith et al., 2008; Hastie and Fithian, 2013; Phillips and Elith, 2013; Wang and Stone, 2019; Ward et al., 2009). In Wang and Stone (2019), we proposed the Constrained LK (CLK) method, which also requires the population prevalence as the prior information. In detail, the CLK method maximizes the following (LK type of) likelihood function,

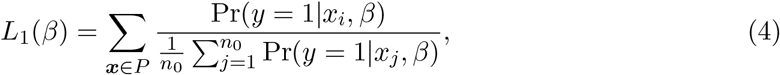

with the constraint, i.e., 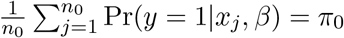, where *π*_0_ is a pre-determined population prevalence and *n*_0_ is the number of background samples. Note that this is different from just maximising the function of Eqn. 3 in the LK method, since the constraint reduces the effective parameter space over which the maximization is performed. It was also illustrated in Wang and Stone (2019) that the CLK method is capable of estimating the true probability of presence, the same as the EM (Ward et al., 2009), SB (Phillips and Elith, 2011) and SC (Steinberg and Cardell, 1992) methods, when the population prevalence is known. However, the population prevalence, *π*, is difficult to obtain in practice.

### 2.3 Estimating Prevalence

Although it is difficult to obtain an accurate pre-determined prevalence estimate of *π* in practice, we find the concept of ‘prototypical presence locations’ proposed by Li et al. (2011) often yields accurate estimates of *π*. The concept was inspired by the presence and background learning (PBL) (Elkan and Noto, 2008), a ML algorithm, and was further developed by Li et al. (2011) in the ecological context of modelling species distributions. In this section, we propose an estimate for the population prevalence *π*, which can be plugged into the CLK method or other methods that require a pre-determined prevalence to ultimately estimate the absolute probability of presence. The steps of the new refined CLK method are summarised in Fig. 1.

**Figure 1:**
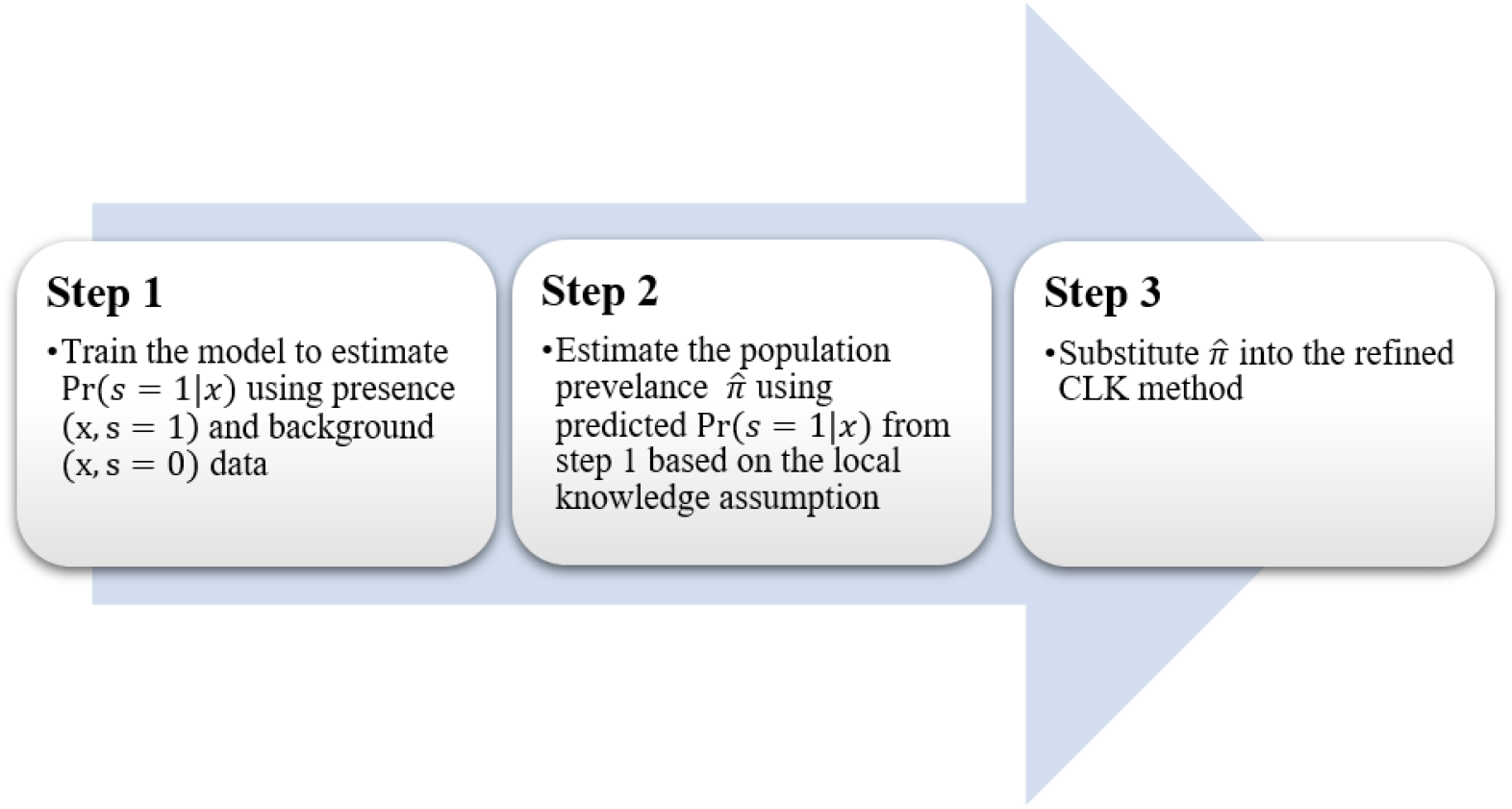
The steps followed in the new refined CLK method.

As a first step, we derive the relation between the probability of a species being observed, i.e., Pr(*s* = 1|*x*), and absolute probability of presence Pr(*y* = 1|*x*) as follows (see derivation in Appendix A),

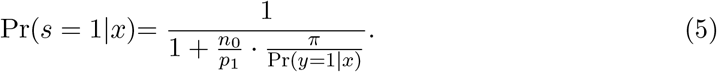

This relationship was also derived by Phillips et al. (2009) and Li et al. (2011). The derivation in Phillips et al. (2009) led to the introduction of the scaled binomial (SB) method (Elith et al., 2011), which assumes the population prevalence *π* to be known in advance.

Although not specifically defined in their work, Elkan and Noto (2008) assume that Pr(*y* = 1|*x*) must equal to 1 at certain *x* values; this property was defined later by Bekker and Davis (2018) as the “local certainty” assumption. Similarly, Li et al. (2011) introduced the concept of “prototypical presence locations” at which the habitat is maximally suitable for the given species to survive. In the statistical language, the conditional probability of presence at these “prototypical presence locations” is one, i.e., Pr(*y* = 1|*x*) = 1. In our approach, a similar assumption will be used to obtain an estimate for *π*.

At prototypical presence locations, i.e. when Pr(*y* = 1|*x*) = 1, it can be shown from 5 that;

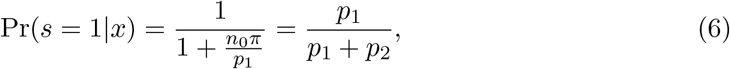

where *p*_1_ is the number of observed presences and *p*_2_ is the number of true presences in the *n*_0_ background data. The ratio of 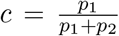 is the probability of a presence point being observed or labelled, i.e., Pr(*s* = 1|*y* = 1).

In another word, Eqn. 6 shows that we can estimate *c* = Pr(*s* = 1|*y* = 1) through the predicted values of Pr(*s* = 1|*x*) at the prototypical presence locations. For this, we use machine learning methods to predict Pr(*s* = 1|*x*), by training a binary classification model using the presence (*x*, *s* = 1) and background (*x*, *s* = 0) data. Popular classification methods, such as logistic regression, k-nearest neighbours (KNN), support vector machines and neural networks, can be used to model Pr(*s* = 1|*x*). In this paper a neural network has been used as a classifier which has been shown to perform very well.

One or more geographical locations should exist as the “prototypical presence locations” but proper identification may be hindered by noise in the predicted values of Pr(*s* = 1|*x*). In order to reduce the effect of noise, Elkan and Noto (2008) proposed three methods for estimating the sampling probability *c*, i.e., the average and the maximum values of Pr(*s* = 1|*x*) at the prototypical presence locations, and the ratio between the sum of Pr(*s* = 1|*x*) at the prototypical presence locations and the sum of Pr(*s* = 1|*x*) over the background. Using the average as an example, *c* can be estimated as follows (Elkan and Noto, 2008; Li et al., 2011).

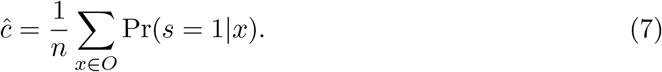

That is, *c* will be evaluated as the mean of predicted probabilities of Pr(*s* = 1|*x*) for *x* that belongs to the prototypical presence locations. According to Eqn. 5, the probability of Pr(*y* = 1|*x*) is an increasing function of Pr(*s* = 1|*x*), so we use the locations where Pr(*s* = 1|*x*) are maximal as the prototypical presence locations. We rank the predicted Pr(*s* = 1|*x*) values for all presence and background points, and those locations where predicted Pr(*s* = 1|*x*) are high are used as the prototypical locations (Li et al., 2011). In our simulation study, we used the top 10th percentile, which is tested sufficient and appropriate for our study.

From Eqn. 6, we can easily show the relation between *π* and *c* as follows;

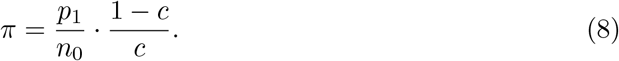

*π* can thus be estimated as,

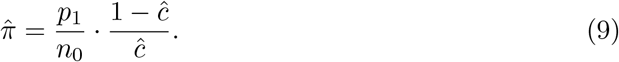

We then substituted 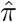 into Eqn. 4 as the constraint of the CLK likelihood function to ultimately estimate the unknown parametric function of Pr(*y* = 1|*x, β*).

In our proposed method, the population prevalence is estimated from the available presence and background data, provided the “local certainty” condition is satisfied. This contrasts with previous studies in which a pre-determined value of prevalence (although often unavailable) is required (Ward et al., 2009; Phillips and Elith, 2011; Wang and Stone, 2019). Population prevalence is the average probability of presence over the whole landscape, and its calculation requires presence information in the whole study area. Compared to the population prevalence, the “local certainty” or the “prototypical presence location” is less constrained that only requires the maximal probability to be one at some sites. This might appear to be a strong assumption, however, we suggest that this assumption is not implausible both theoretically and practically.

### 2.4 Justification of Local Certainty/Prototypical Presence Location

In the relevant ML literature such as Elkan and Noto (2008) and Li et al. (2011), it is typically assumed that the “prototypical presence location” or the “local certainty” condition holds for general classification problems. However, these critical assumptions were not made explicit in the literature. In this section, we will explain how the “prototypical presence location” or the “local certainty” is not implausible in the context of species niche modelling. To understand the concept clearly, consider the problem of classification when working with presence-absence data. Suppose for example, a logistic function for Pr(*y* = 1|*x*) (such as in Eqn. 1) is the underlying resource selection probability function that drives the distribution of presence and absence observations. What we do in simulating the data is to firstly generate a random value *r* on the interval [0, 1] for each pixel. A pixel is assigned *y* = 1 (presence) if *r* ≤ Pr(*y* = 1|*x*) and *y* = 0 (absence) otherwise. Note that a pixel with a probability less than one could be randomly assigned as either presence or absence (or just absence) depending on the value of *r*. In order to have a unique region in the feature space that is only occupied by the presence samples, some pixels must have Pr(*y* = 1|*x*) = 1 (“prototypical presence locations”). Otherwise wherever a presence point is observed in the feature space, it would mix with an absence point and it will be hard to distinguish the two classes. This would render the classification impossible.

Fig. 2 shows two types of presence-absence data. The data points (red for presence and black for absence) on the left plot of Fig. 2 were generated from a logistic probability distribution function that satisfies the local certainty condition. It can be seen that there are some “prototypical presence” points (red) that are clearly separated from the absence (black) points especially in the top left quadrant. On the right side of Fig. 2, data points were generated from a scaled logistic function, where the maximal probability does not reach one. In contrast to the left plot, both presence and absence points nearly overlap in the feature space of (*x,y*), and there appears no “prototypical presences” region where presences are clearly separated from absences. ML focuses more on classes that are strongly separable, and the two classes in the second example would be hard to separate, regardless of method being used. Therefore we believe that the local certainty or the prototypical presence location holds for general classification problems in the context of ML, so that the two classes (presence and absence) are “separable” in feature space.

**Figure 2:**
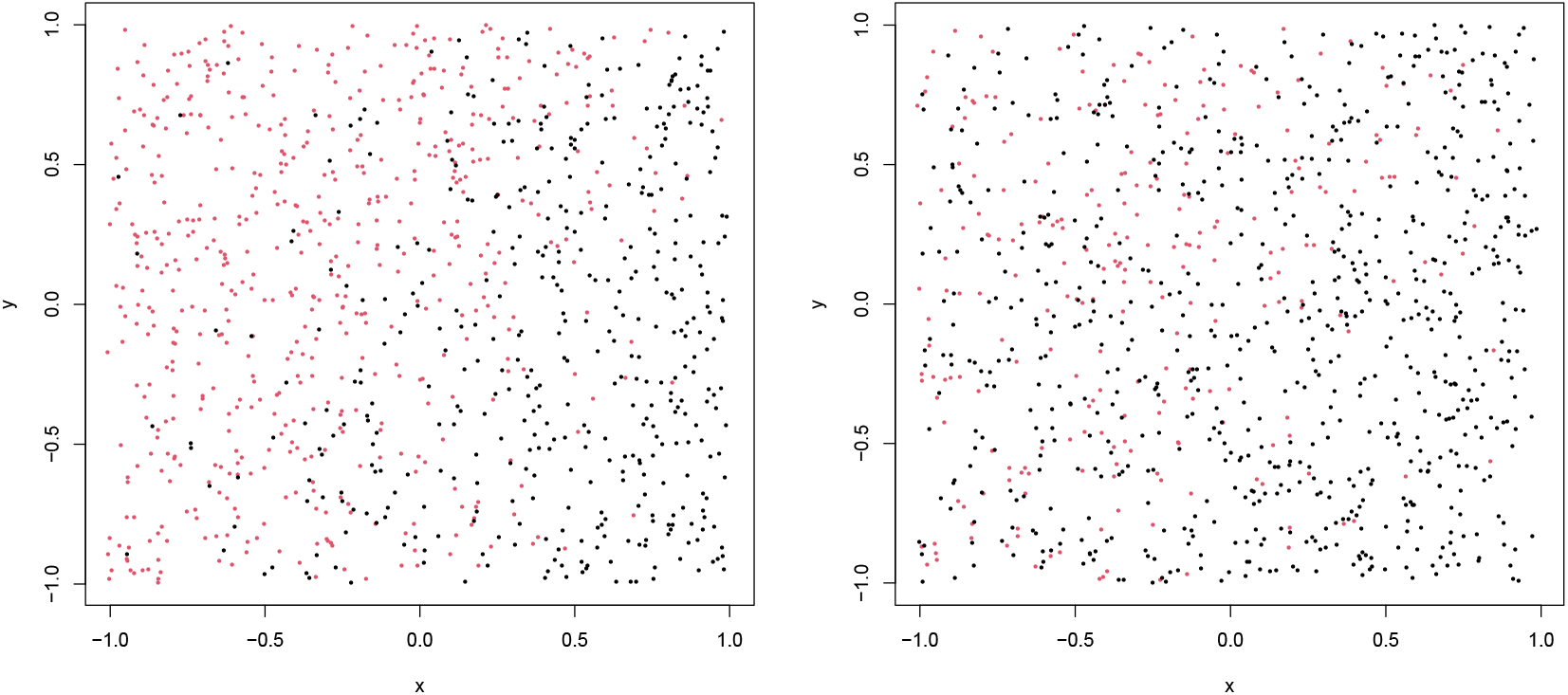
The left plot shows the two classes, presence (red dots) and absence (black dots), generated from a logistic probability function 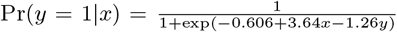, which satisfies the local certainty condition, i.e., Pr(*y* = 1|*x*) = 1 at some points. The right panel shows the two classes generated from a scaled logistic probability function 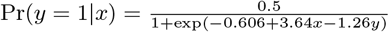, where the probability of one is not reached in the feature space of (*x,y*). It is obvious that the presence (red) and absences (black) points are more separable on the left, whereas the two classes overlap and hard to separate in the right plot.

From a practical perspective, local certainty holds, for example, if some locations or combination of niches always provide sustainable conditions for a species to survive. This should be satisfied for an ecosystems in an equilibrium state, where there exists sustainable conditions suitable for species to survive at least in some areas.

### 2.5 Local Knowledge Condition

In statistical experiments, however, we notice that the local certainty or prototypical condition is sometimes not satisfied, i.e., Pr(*y* = 1|*x*) does not reach the maximal value of one. The scaled logistic example used in Hastie and Fithian (2013) to demonstrate the failure of the LK method as well as our second example in Fig. 2 are such examples. In this circumstance when the classes of presence and absence may not be distinguishable (in the context of ML), we believe extra assumption or condition must be required to identify probability of presence from presence-background data. However, this condition should not be based on unfounded assumption, such as the RSPF conditions (Solymos and Lele, 2015). Rather in this paper, we propose a “local knowledge” condition that is one of such assumptions or conditions to handle these “difficult” problems. Compared to the commonly recognised population prevalence, which requires the probabilities of presence over the whole landscape, the local knowledge condition relaxes the information required and is thus less constrained.

A simple derivation from Eqn. 5 shows that;

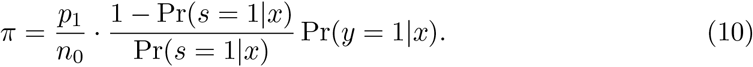

If we have the knowledge, for example, about the maximal probability of presence of the species (which is not necessarily one), we can rank the predicted values of Pr(*s* = 1]*x*) for all observed presences and background points. Those locations with the highest Pr(*s* = 1|*x*) will be used as the locations with ”local knowledge”. Using a maximal probability of 0.7 as an example, we can show from Eqn. 10 that

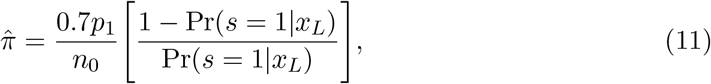

where *L* are those locations with local knowledge, i.e., the highest probabilities of Pr(*s* = 1|*x*) in this case. Local knowledge does not necessarily refer to the information of the maximal probability, and it can also include any nominated probability of presence at a given site or over a particular landscape. In general, if “local knowledge” is available so that there are some locations where the probabilities of Pr(*y* = 1|*x*) are known, it should be possible to make use of the trained value of Pr(*s* = 1|*x*) at these locations to estimate *π* from Eqn. 10 and to finally estimate our objective function of Pr(*y* = 1|*x*). In the following simulation section, the scaled logistic example explored in Hastie and Fithian (2013) will be revisited to show how the local knowledge assumption can be used to estimate the absolute probability of presence.

The local knowledge condition was built under the same framework and extends the local certainty condition. The “prototypical presence location” assumption given by Elkan and Noto (2008) and Li et al. (2011) is just one particular “local knowledge” subclass, in this case where Pr(*y* = 1|*x*) = 1 at these locations. In practice, local knowledge could be gained through experts’ knowledge or pilot studies.

At the end of this section, the steps (as given in Fig. 1) of the new refined CLK method to estimate the conditional probability of presence Pr(*y* = 1|*x, β*), is summarized as follows:

Step 1) Train the model to estimate Pr(*s* = 1|*x*), using all presence (*x*, *s* = 1) and background (*x*, *s* = 0) data points. A commonly used ML binary classifier, such as the neural network or support vector machine, can be used for modelling Pr(*s* = 1|*x*);

Step 2) Estimate the population prevalence 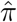 using Eqn. 10 based on the predicted probabilities of Pr(*s* = 1|*x*) at the prototypical presences locations or locations with local knowledge;

Step 3) Maximize the log-likelihood function of 4 with respect to *β*, with *π* substituted by 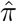 in step 3 and obtain the estimated probability of presence Pr(*y* = 1|*x*; *β*).

### 2.6 Simulation Study

In this section, simulations are carried out firstly to assess the fragility of the LK method and its RSPF conditions proposed by Lele and Keim (2006) and Solymos and Lele (2015); and secondly to demonstrate the performance of the proposed concept of “local knowledge”, in conjunction with the CLK method, to estimate the probability of presence.

To simplify the presentation, we consider a single predictor variable *x* in our simulations. We use three different groups of functions for Pr(*y* = 1|*x*). Category 1 meets both the RSPF and local certainty (LC) conditions (see the top plot of Fig. 3 and top section of Table 1); Category 2 functions only satisfy the local certainty condition (see the middle plot of Fig. 3 and middle section in Table 1) (by this we mean that the selected parametric functions achieves the probability of 1 at some points of *x*) and Category 3 functions only satisfy the RSPF conditions and not the local certainty (see the bottom plot of Fig. 3 and bottom section in Table 1). In each category, three functions are selected such that it consist of one linear, one quadratic and one cubic function of *x* (see Fig. 3 and Table 1). To ensure functions in Category 1 and Category 3 satisfy the RSPF conditions, we used exactly the same functions as those in Solymos and Lele (2015) that include linear, quadratic and cubic logistic and complementary log-log (Cloglog) functions. In Category 2, linear scaled logistic, Gaussian and cubic exponential functions are chosen as not to satisfy the RSPF conditions, because the constant terms in these functions would be cancelled out in the LK’s loglikelihood function (see Eqn. 3). Both the scaled logistic and exponential functions were also noted in Solymos and Lele (2015) as not satisfying the RSPF conditions. These functions have been modified in our study to meet the local certainty conditions (see Fig. 5 upper row).

**Figure 3:**
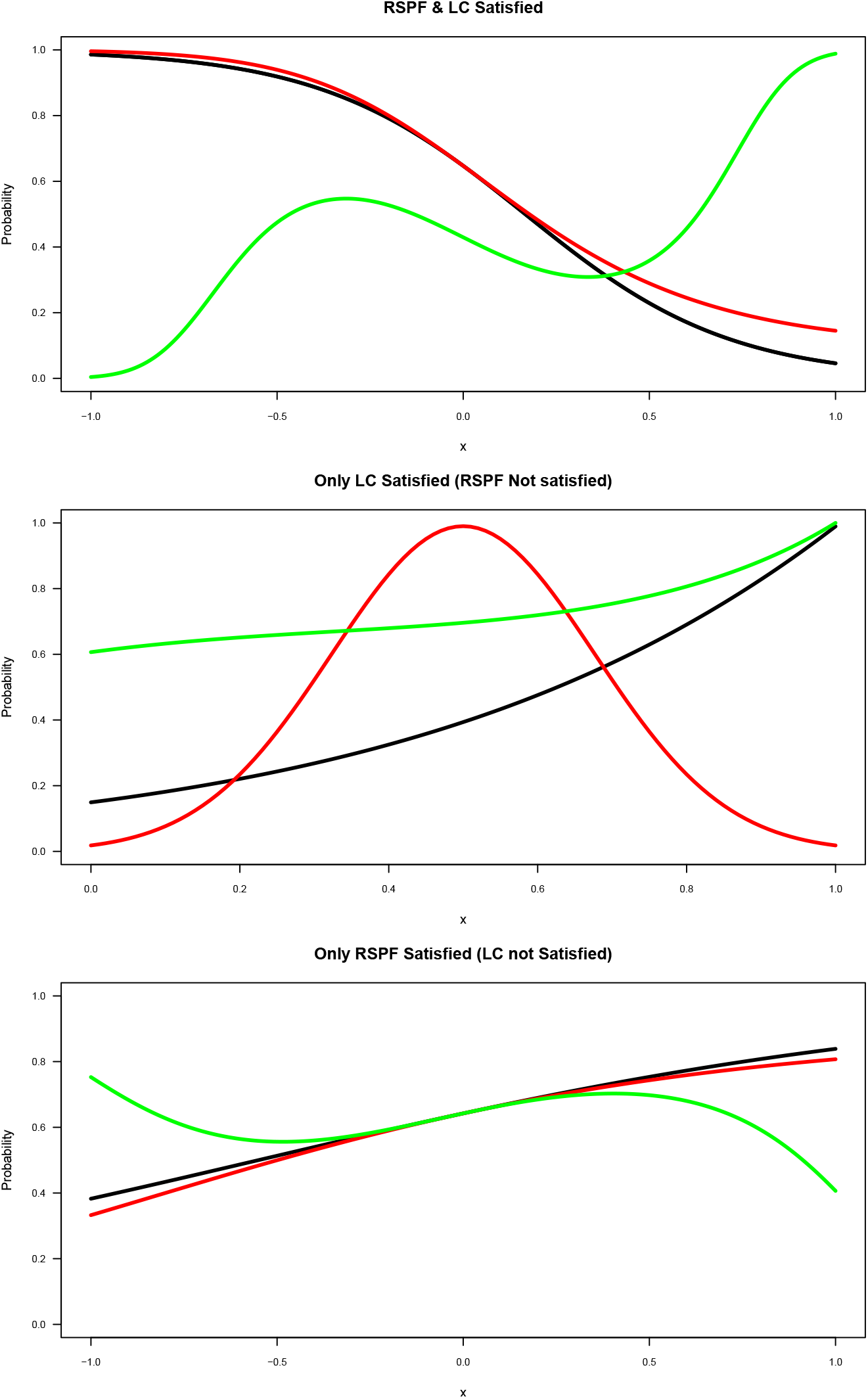
The selected simulated species distributions of the three categories of functions. The Pr(*y* = 1|*x*) is plotted as a function of the covariate *x*. The top plot shows Category 1 functions (both RSPF and LC satisfied) which include linear logistic (black), quadratic logistic (red) and cubic logistic (green). The middle plot gives functions of Category 2 (only local certainty satisfied/RSPF not satisfied) which are Linear Scaled logistic (black), Gaussian (quadratic) (red) and cubic exponential (green) functions. The bottom plot shows Category 3 (only RSPF satisfied) which are linear logistic (black), quadratic logistic (red) and cubic logistic (green) functions.

**Table 1:** Probability of presence for the simulated species used in the experimental evaluation, where Category 1 satisfies both RSPF and LC conditions, Category 2 is where only LC is satisfied (RSPF conditions are not satisfied). Category 3 is where only RSPF conditions are satisfied (LC is not satisfied).

| Category | Conditions Satisfied | Function Type | Probability Functions |
| --- | --- | --- | --- |
| Category 1 | Local | Linear | $\Pr(y = 1\vert x) = \frac{1}{1+\exp(-0.606+3.64x)}$ |
| | Certainty & | Quadratic | $\Pr(y = 1\vert x) = \frac{1}{1+\exp(-0.606+3.64x-1.26x^2)}$ |
| | RSPF | Cubic | $\Pr(y = 1\vert x) = \frac{1}{1+\exp(0.284+2.298x+0.232x^2-7.269x^3)}$ |
| Category 2 | Local | Scaled logistic | $\Pr(y = 1\vert x) = \frac{8.3}{1+\exp(4-2x)}$ |
| | Certainty | Gaussian | $\Pr(y = 1\vert x) = 0.99 \exp(-(4x - 2)^2)$ |
| | | Exponential | $\Pr(y = 1\vert x) = \exp(-0.5 + 0.5x - 0.9x^2 + 0.9x^3)$ |
| Category 3 | RSPF | Linear | $\Pr(y = 1\vert x) = \frac{1}{1+\exp(-0.5855-1.064x)}$ |
| | | Quadratic | $\Pr(y = 1\vert x) = \frac{1}{1+\exp(-0.5855-1.064x+0.218x^2)}$ |
| | | Cubic | $\Pr(y = 1\vert x) = \frac{1}{1+\exp(-0.5855-1.064x+0.218x^2+1.81x^3)}$ |

All the functions in Categories 1 and 3 are fitted by logistic functions. Gaussian and cubic exponential functions in Category 2 are fitted by exponential functions to account for the format of the original functions. The scaled logistic function in Category 2 is also fit by an exponential function to test the robustness of the CLK method with different type of fitting function than the logistic function (See Fig. 5 top row). For Category 3, the maximum probabilities of 0.83, 0.8 and 0.75 are used as the local knowledge conditions for the linear, quadratic and cubic logistic functions respectively. We also test the performance of the CLK method with the mis-specified local knowledge information, where the local knowledge used differs the true probability by ±10%. Results are shown in the bottom row of Fig. 5.

The Cloglog functions studied by Solymos and Lele (2015) are also investigated to assess the performance of the methods when the fitting models are logistic parametric functions. The results are shown in Figs C.2 and C.3 in appendices. For Category 2 functions in cloglog simulation, where only RSPF conditions are satisfied, the maximal probabilities of 0.73, 0.89 and 0.68 are used separately as the local knowledge conditions for linear, quadratic and cubic functions (see Fig. C.3).

Solymos and Lele (2015) pointed out that if the non-linearity on the log-scale is weak, it may need very large sample sizes to get reasonable estimate for the LK method. In order to ensure a stable performance of the LK method, the simulations are carried out with 5,000 presence samples and 50,000 background samples. When the LK method results are not satisfactory in some experiments such as Category 3 simulations, the experiments are repeated with even larger samples of 50, 000 presences and 500, 000 background points, in order to investigate the performance of LK method under very large sample conditions. (Phillips and Elith, 2013; Solymos and Lele, 2015). We also run simulations with a small sample, 250 presences and 5,000 background points for functions in Category 1 to test the robustness of the LK and CLK method. The small sample results are given in Fig. 4 bottom row.

**Figure 4:**
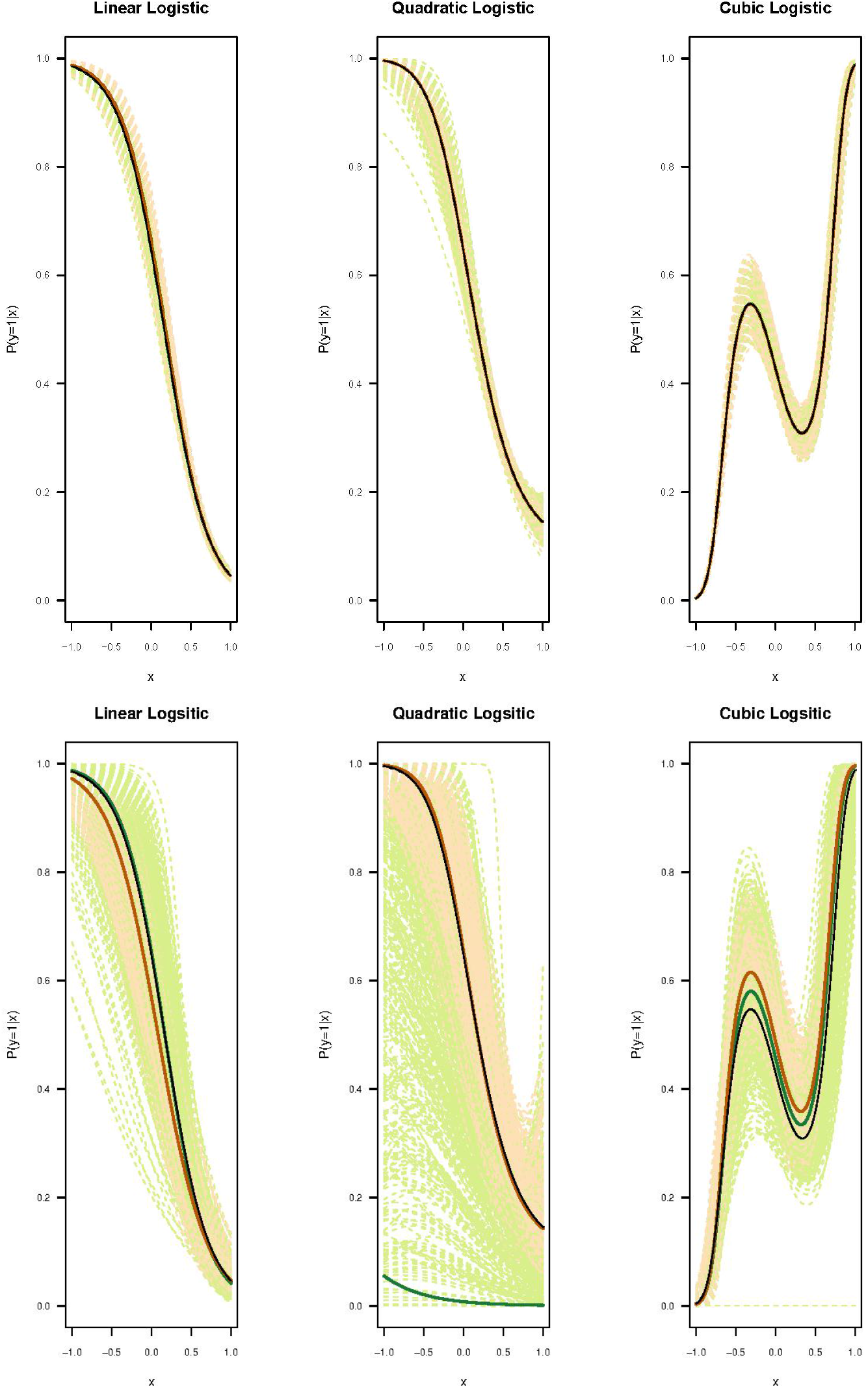
Top panel contains the graphs of Category 1, in which the Pr(*y* = 1|*x*) is plotted against the covariate *x* when the original parametric function satisfies both the RSPF and local certainty conditions. For each simulation, 5, 000 presence samples and 50, 000 background samples were chosen from a landscape with the predictor variable *x* in [−1, 1]. The bottom panel contains the category 1 functions with a smaller sample of 250 presence samples and 5000 background samples chosen from the same landscape as the top panel. Estimated lines from each of 1000 simulations are plotted in light green (LK) and light orange (CLK). Three solid lines are plotted representing the true probability function (black), representing the average LK estimates (green) and average CLK estimates (orange) respectively, as averaged from 1000 simulations.

The neural network package ‘nnet’ in R (Venables and Ripley, 2002) (https://CRAN.R-project.org/package=nnet) was used to train Pr(*s* = 1|*x*) for the refined CLK method. We rank the predicted Pr(*s* = 1]*x*) for the presence and background points, and those locations whose predicted Pr(*s* = 1|*x*) lie in the top 10th percentile are used as the prototypical locations or locations with the local knowledge. The choice for the 10th percentile threshold is arbitrary but was tested sufficient and robust for our simulation studies.

We run the simulations 1000 times for each species distribution function. The estimated environmental relationship for each of the 1000 simulation are plotted using the LK and refined CLK methods, against the true probabilities. Root mean square errors (RMSE) of fitted probabilities against the true probabilities of presence are also calculated and plotted in Figs. 6, C.2 and C.3. Moreover, the error (non-convergence) rates of the LK method in 1000 iterations are given in Tables D. 1 and D. 2 in appendix.

All model fitting and assessment have been carried out in R version 3.6.1. The R code supplied by Solymos and Lele (2015) has been used to choose the functions that satisfy the RSPF condition. The ‘ResourceSelection’ package (Lele and Keim, 2006; Lele, 2009; Solymos and Lele, 2015) has been used to fit the LK method, and the R code provided by Phillips and Elith (2013) has been used with modification where relevant.

### 2.7 Revisiting an Example Hastie and Fithian (2013)

One of the pivotal publications by Hastie and Fithian (2013) demonstrated the failure of the LK method to fit and distinguish between the scaled and full logistic functions. From the perspective of ML, the classes of presence and absence generated from a scaled logistic function are actually not separable, because scaled logistic functions do not satisfy the local certainty condition. We revisit the example in Hastie and Fithian (2013) and fit the scaled logistic function with the Constrained LK method but using the “local knowledge” information. We will show that the “local knowledge” assumption is not only helpful but also necessary in removing the multiple solutions that create identifiability issues in the LK method.

In their paper, Hastie and Fithian (2013) demonstrated the lack of fit in the LK methods using two logistic functions. The full logistic function considered is 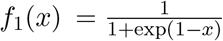 while the scaled logistic function is exactly half of that, i.e., 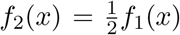, where *x* ∈ [−2.5, 2.5]. We first fit both functions by the LK and CLK methods, using a sample of 5,000 presences and 50,000 background data points with 1,000 iterations (top row of Fig. 7). We also tried different sample size to test the robustness of the LK method as well as the CLK method (see bottom row of Fig. 7). For the refined CLK method, we assumed the local knowledge of Pr(*y* = 1|*x*) = 0.8 and Pr(*y* = 1|*x*) = 0.4, which are the maximal probabilities that the full and scaled logistic functions can reach over the range of *x*. We used the top 10th percentile of sites where Pr(*s* = 1|*x*) are highest as the locations with local knowledge.

## 3 Results

### 3.1 Simulation Results

The results of the three categories are given in Fig. 4 & 5 in which the Pr(*y* = 1|*x*) is plotted against *x*. In these figures, the black line gives the original parametric functions selected as given in Table 1. The green and orange dashed lines indicate the simulation lines for the 1000 iterations carried out for the LK and refined CLK methods while the respective solid lines show the average of those simulations. The top panel of Fig. 4 shows that when both the RSPF and the local certainty conditions are satisfied in Category 1, the LK and refined CLK methods both perform well in estimating the true function of Pr(*y* = 1|*x*), with the mean estimates close to the true functions. Both methods have similarly small RMSEs for the fitted probabilities against the true probabilities (see top left panel of Fig. 6). However, when the number of presence points reduces (such as to 250), the LK method start to perform more unstably with much more variations (see Fig. 4 bottom row). The dispersion for the CLK method also increase, but they are consistently stable. The CLK method with the local certainty condition still perform well with small samples regarding both the accuracy and precision (see top right plot in Fig. 6).

**Figure 5:**
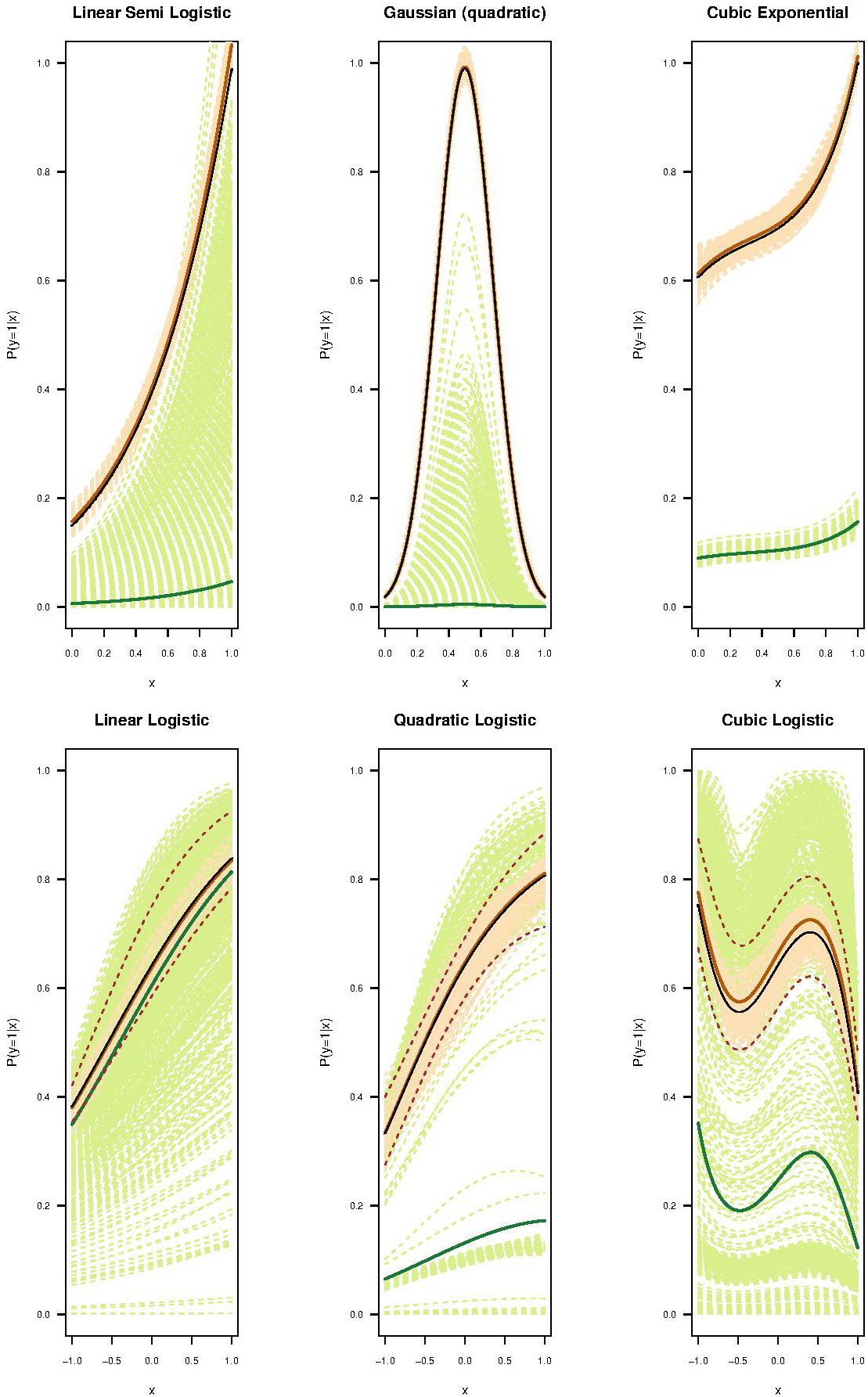
Category 3: The top panel gives the graphs of Category 2 in which RSPF is not satisfied; only LC is satisfied. In the bottom panel Pr(*y* = 1|*x*) is plotted as a function of *x* when the original logistic parametric function only satisfies the RSPF condition with local knowledge of of the maximal probabilities of 0.83 for the linear logistic (left), 0.8 for Quadratic (middle) and 0.75 for Cubic (right) logistic functions. Estimated lines from each of 1000 simulations are plotted in light green (LK) and light orange (CLK). Three solid lines are plotted representing the true probability function (black), the average LK estimates (green) and average CLK estimates (orange) respectively, as averaged from 1000 simulations. The dashed brown lines in the bottom row indicate a single run of the simulation when the local knowledge is misspecified by ±10% from its true value. For each simulation, 5, 000 presence samples and 50, 000 background samples were chosen from a landscape with the predictor variable *x* in [−1, 1].

**Figure 6:**
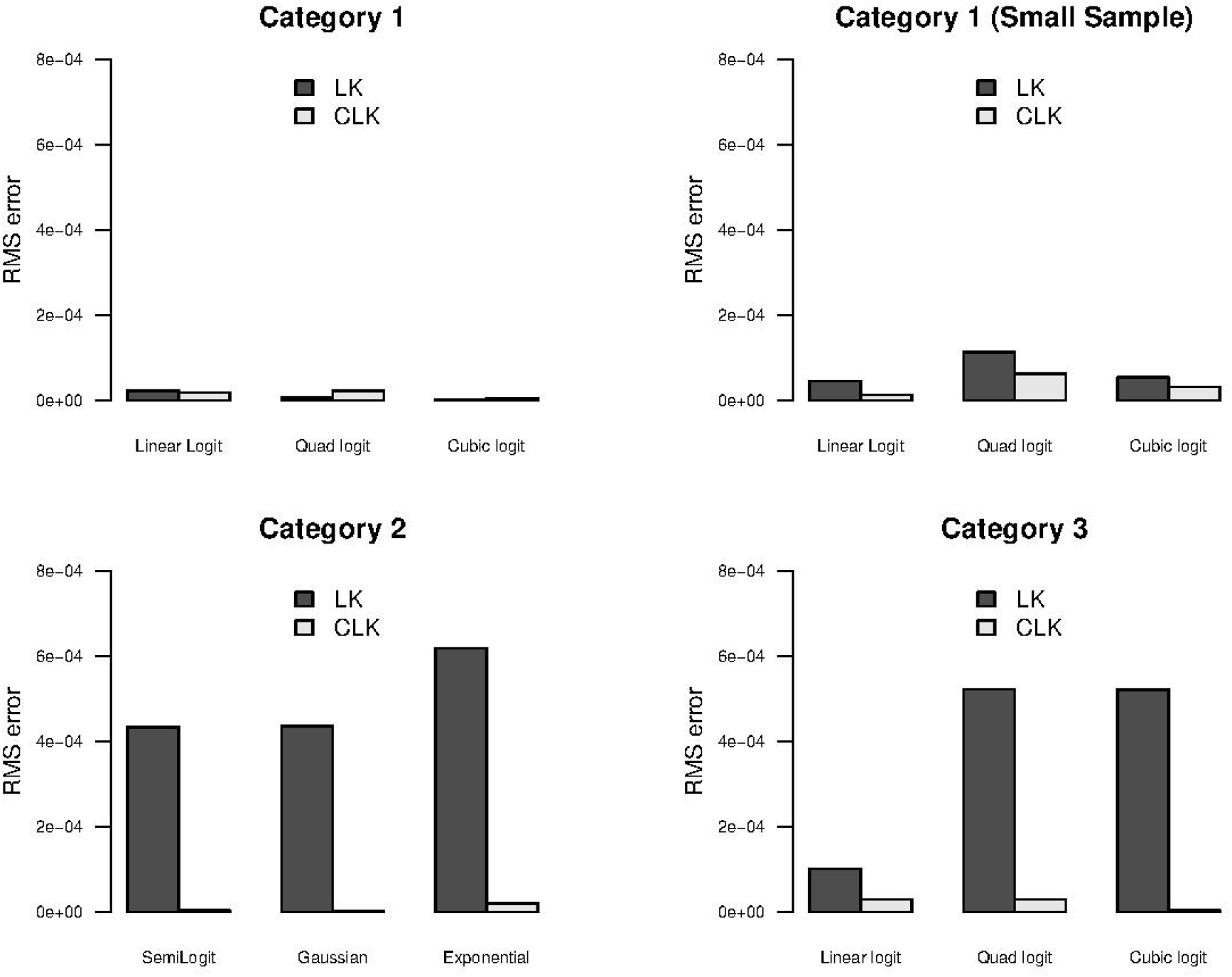
The bar plot of the RMSEs of the two methods, LK and CLK for the logistic functions in Category 1 (top left) as given in top row in Fig. 4, Category 1 (top right) with small samples, Category 2 (bottom left) as given in top row in Fig. 5 and Category 3 with local knowledge of 0.7 (bottom right) as given in the bottom row of Fig. 5.

Category 2 functions (Fig. 4 bottom panel) do not satisfy the RSPF conditions, but they meet the local certainty conditions. It means that the selected parametric functions achieve the probability of 1 at some points of *x*. Under these circumstances, the refined CLK method performs consistently well for all functions (orange), while the LK (green) performs poorly as expected given that these functions fail to satisfy the RSPF conditions. The poor performance is seen by the large variability and the simulated curves not being even close to the original function (cubic exponential function) in the LK simulations that indicate an identifiability issue. The RMSEs (Fig. 6 bottom left panel) clearly display the poor performance of the LK method with large RMSEs and the better performance of the refined CLK method (relatively smaller RMSEs).

Fig. 5 bottom panel shows the performance of the two methods for Category 3, when they both satisfy only the RSPF conditions. Interestingly, the LK method (green lines) fails to perform well in all functions in Category 3, where the claimed RSPF conditions are all met for these functions (Solymos and Lele, 2015). The average fit for the linear logistic function may appear close to the true probability function, but the individual estimates are so unstable that fluctuate widely. On the other hand, the refined CLK method well predicts the true resource selection probability function when accurate local knowledge information is available. The bottom right-side panel of Fig. 6 shows the large RMSEs of the LK method for all three functions in Category 3.

The brown dashed lines in the bottom panel of Fig. 5 show the estimates of probability when the local knowledge is mis-specified by a ±10% bias from the true local knowledge. The mis-specification of the local knowledge condition leads to limited but consistent over or under-estimation of the probability distribution functions. Regardless, the misspecified estimates of the CLK method still outperform the LK method in all cases. Again partially due to more information utilised, the CLK estimates have significantly small variances compared to those of the LK method. The Cloglog functions that are fitted using logistic parametric functions confirm the superior performance of the refined CLK method in terms of both accuracy and dispersion when fitting the model with mis-specified parametric functions (see Figs. C.2 and C.3 in Appendix).

Phillips and Elith (2013) and Solymos and Lele (2015) mention that the LK method performs better when the sample size is ‘large enough’. Solymos and Lele (2015) hint that the above-mentioned problem can be overcome with larger sample sizes. We found that when using a ‘very large’ sample of 50,000 presences and 500,000 background samples (see Fig. B.1 in Appendix), the LK method shows an improved performance with less variations when compared with the performance from a sample of 5, 000 presences and 50, 000 background samples (See Fig. 5). Nevertheless, a significant improvement in performance is only observed for some functions (linear logistic function in Fig. C.1 in Appendix), while for other functions the improvement is only slight.

An instability of the LK method is also observed while using the ‘ResourceSelection’ package in fitting the LK method. The LK shows a greater variation in the estimation result, when different starting values are used to optimize the log-likelihood function. Even in the cases when the logistic parametric functions satisfy the RSPF conditions, failures (non-convergences) are observed in the simulations. The non-convergence rate (see Table D. 1 in Appendices) show the proportion of simulations that don’t converge. Using the quadratic logistic function in category 1 as an example, the LK method didn’t converge 2.2% of times when the function satisfies the local certainty condition. The failure rate increases to 31.4% (for the quadratic logistic function in category 3) when the local certainty is not met, while in both cases the quadratic functions still satisfy the RSPF condition. Similar behaviour is also observed for mis-specified models (See Table D. 2 in the Appendices).

### 3.2 Results of the Revisited Example from Hastie and Fithian (2013)

Hastie and Fithian (2013) showed that the proportional likelihood (see Eqn. 2) that forms the basis of the LK method has identifiability issues. In their argument they considered the full logistic function *f*_1_(*x*) and a scaled logistic *f*_2_(*x*) which is exactly half of the full logistic function *f*_1_(*x*). The top panels in Fig. 7, plot these two functions *f*_1_(*x*) & *f*_2_(*x*) given by Hastie and Fithian (2013).

**Figure 7:**
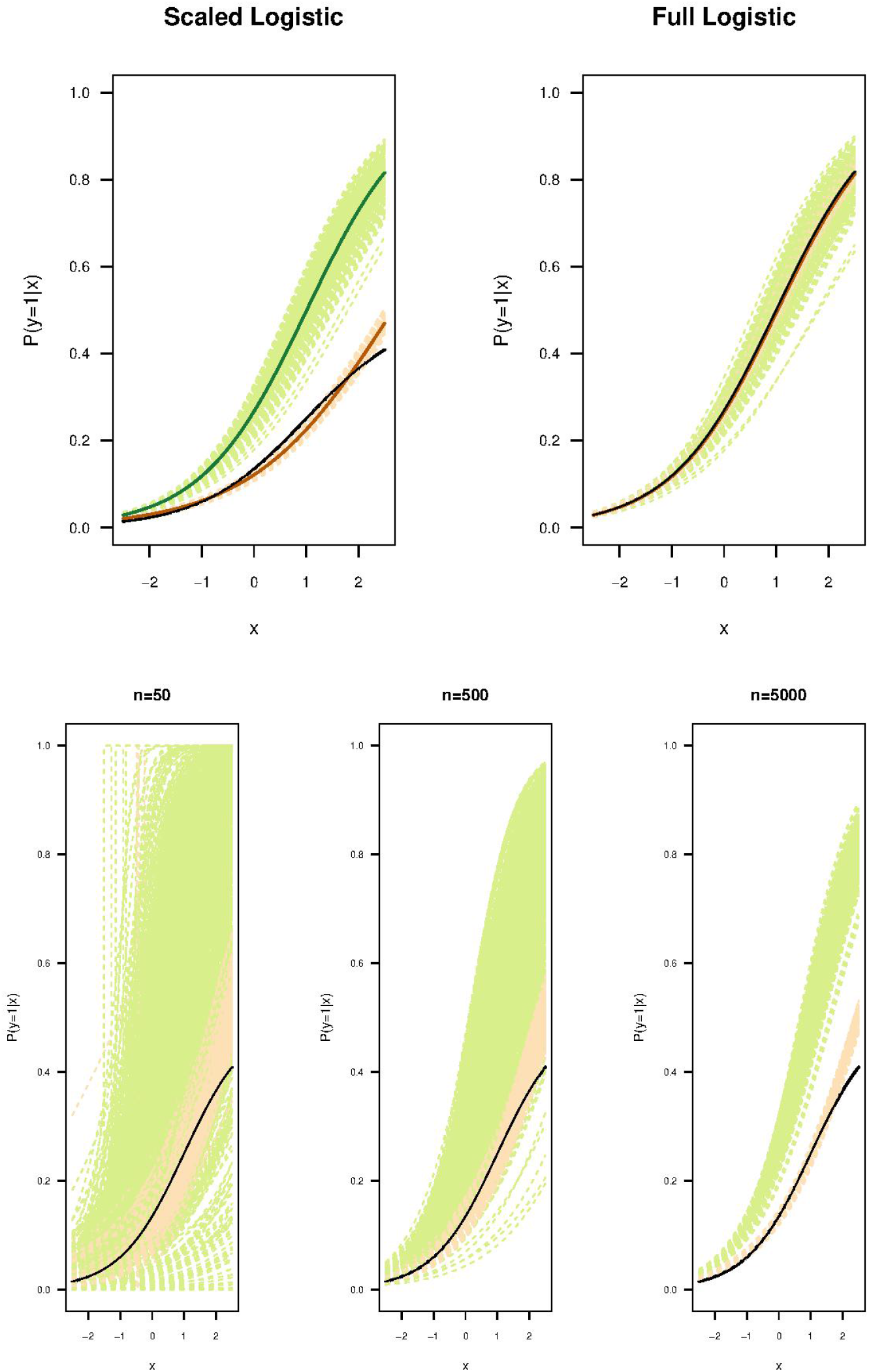
Top panel shows the full and scaled logistic functions (same as Hastie and Fithian (2013)). Here we used the ‘local knowledge’ of Pr(*y* = 1|*x*) = 0.8 for the full logistic function and Pr(*y* = 1|*x*) = 0.4 for the scaled logistic function both around *x* = 2.5. The green and orange dashed lines (top panel) represent estimates of LK and CLK methods respectively and the solid green and orange lines represent the averages of the 1000 simulations. The original logistic functions are given in black. The bottom panel shows the fitted scaled logistic functions from both the LK method (green) and the CLK (orange) with different sample sizes. From the bottom left to right, the presence and background samples increase from (50, 5000), (500, 5000) to a very large samples of (5000, 50000).

Because of the scaling, it is easy to see from Eqn. 3 that the likelihoods of these two functions *f*_1_(*x*) and *f*_2_(*x*) are exactly the same. Thus, when we attempted to fit the two different datasets (generated from the scaled and the full logistic function), the LK method gave exactly the same parameter estimates for the *β_i_* for both functions. This is confirmed in our simulation when large sample sizes are applied. The LK method yields the same estimates for both the full and the scaled logistic functions. Thus despite the fact that 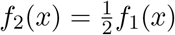 (black line in Fig. 7 top left panel), the estimation scheme finds 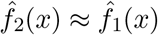. Fig. 7 (top left panel) shows the LK fits (green lines) are close to the full-logistic (plotted in black Fig. 7 top right panel). This effect was observed and explained in Hastie and Fithian (2013).

The refined CLK method, on the other hand, predicts both logistic functions very well, when the maximal probabilities for Pr(*y* = 1|*x*) are available for both the full and scaled logistic regressions. The predicted functions from the refined CLK method align closely with both the true probability functions, with small variations observed between replications (see orange dashed lines in the top panel of Fig. 7). This example shows that local knowledge assumption is not just useful but is necessary to retrieve the scaled logistic function, with which the LK method has failed, when the local certainty condition is not satisfied.

An effect that wasn’t discussed in Hastie and Fithian (2013) concerns the case when the sample sizes of the presence and/or background are small (bottom panel of Fig. 7). Now the identifiability problems of the LK method are even more severe. The green dashed lines show that the many LK-fits we attempted are in fact a manifestation of the likelihood’s multiple proportional solutions interfering. Each such solution has the same likelihood as the full logistic model. As the sample sizes increase the model gradually identifies the function with the ‘maximum’ likelihood as the interested parametric function, which in these types of cases tends to be the full logistic function. This is why the LK method has such trouble fitting scaled functions. Moreover, we also found that it requires very large samples for the LK method to perform stably (bottom panel of Fig. 7). By contrast, the CLK method performs consistently well even with a small number of presence points.

## 4 Discussion

In this paper, we have studied the modelling of the probability of presence (or probability of use) of a species using presence-background (or use-availability) data. In order to estimate the resource selection probability, either a strong parametric assumption such as the logit function requirement for the LK method or extra information such as the population prevalence (as in the SB, EM and constrained LK methods) has been proposed (Ward et al., 2009; Elith et al., 2011; Phillips and Elith, 2013; Wang and Stone, 2019).

The LK method, along with other methods with a strong parametric assumption, have been widely used in the literature and it has been claimed that the absolute probability of presence may possibly be estimated without requiring an estimate of the prevalence (Lele and Keim, 2006; Solymos and Lele, 2015). But there have been arguments in the literature and many researchers have warned against using these methods to accurately estimate the actual probability of presence (Elith et al., 2006; Guillera-Arroita et al., 2015; Hastie and Fithian, 2013; Keating and Cherry, 2004; Knape and Korner-Nievergelt, 2015; Phillips and Elith, 2013; Renner and Warton, 2013; Wang and Stone, 2019; Ward et al., 2009). Simulation study suggests that the LK method performs well only when the “RSPF-eligible” functions satisfy the local certainty condition and samples are not small. When presence sample is small, linear logit/cloglog functions with the local certainty seem the only functions that performs reasonably well. Taking a large sample along with a strong parametric assumption may sometimes overcome these problems, as we think Solymos and Lele are arguing. But either finding a large enough sample or knowing a parametric assumption is not feasible in practice. Our paper shows again that these methods (such as the LK) are fragile and can give unreliable estimates even when its underlying so-called RSPF condition is met. Thus our results contradict the claim made in Solymos and Lele (2015) that “if the RSPF condition is satisfied, it is possible to estimate absolute probability of selection”. Although the RSPF condition is necessary for the LK method, our study indicates the “success” of the LK method depends crucially on the local certainty, i.e., two classes of presence and absences are separable.

It was shown that the LK method has difficulty distinguishing between the actual likelihood and other possibly similar likelihoods that arise from scaled functions. From the perspective of ML, the presence observations generated from the scaled logistic functions, which do not satisfy the local certainty condition, completely overlap with absence observations and two classes are inseparable. It is thus expected that the LK method would have multiple solutions and difficulty in locating the true probability function of presence. Under this circumstance, we believe that not only the LK method but any other methods should have trouble in distinguishing two inseparable classes, except when extra information is enforced. The extra information, however if coming from unfounded model assumption, only renders fragile estimation/prediction of the desired probabilities. Rather the local knowledge proposed in the paper provides an effective additional datum to reliably estimate absolute (rather than relative) probability of presence from the PB data. This condition further relaxes the commonly recognised population prevalence and helps to overcome the identifiability issue inherent in the proportional LK likelihood function i.e., the problem Hastie and Fithian (2013) elaborated on.

Other methods, such as the SB (Phillips and Elith, 2011), EM (Ward et al., 2009) and CLK (Wang and Stone, 2019), require the population prevalence to estimate the absolute probability of presence from the PB data. Information concerning population prevalence, however, is hard to obtain in practice. We propose a new approach which combines both ML (Elkan and Noto, 2008; Li et al., 2011) and statistical techniques to strengthen the estimation of population prevalence and to finally estimate the probability of presence. Our approach estimates the average probability of presence (population prevalence) by utilising the local certainty assumption, which was justified as not implausible both theoretically and practically. By doing this, we eliminate the strong parametric assumption in the LK method, while removing the need of additional fieldwork to estimate the population prevalence. As a result, our estimate of the population prevalence can also be fed into the SB, EM as well as the widely used MAXENT (Phillips et al., 2006) models to predict the absolute probability of presence.

The local certainty condition assumes that Pr(*y* = 1|*x*) = 1 is satisfied at the prototypical presence locations in a balanced ecosystem. As we describe, users can find prototypical presence locations by identifying large prediction values of Pr(*s* = 1|*x*) from the observed presences and background points (or the presence locations). As a reviewer pointed out, users could enlarge the study area so that it is more likely to include certain prototypical presence locations in the study area. Additionally, adding more relevant features to distinguish between species’ presences and absences may also help because the condition of Pr(*y* = 1|*x*) = 1 at some points should be satisfied for those systems in a sustainable equilibrium state, or when there are sustainable conditions suitable for species to survive at some sites. On the other hand, we believe that it becomes a philosophical discussion as to whether the local certainty should be regarded as an inherent requirement for the classification problem in ML or as a model assumption in the language of statistics. In statistics, methods are governed by model assumption and these regularly allow classes to overlap. Statisticians are interested in robustness which refers to violation of model assumptions. ML on the other hand focuses more on classes that are strongly separated but of complex shapes; therefore ”separable” would be regarded as an inherent requirement otherwise overlap would be an issue regardless of the ML method used (Trappenberg and Back, 2000; Xiong et al., 2010).

The “prototypical presence” (local certainty) condition is not always met especially in statistical modelling. The “local knowledge” condition proposed in the paper is an extension of the “prototypical presence” (local certainty) condition. It serves as a necessary condition to resolve the identifiability issues in those resource selection probability functions that result in “inseparable” classes of presence and absence. Other related ideas of anchoring the presence probability have also been suggested, such as using extra effort at a subset of sites (e.g. with occupancy model designs) or integration of presence-only with presence-absence data (Fithian et al., 2015; Koshkina et al., 2017; Dorazio et al., 2015). The conceptual idea behind all these methods is the same, i.e., using additional information to anchor the presence background data in order to estimate probability of presence. Compared to these extra information, we believe the “local knowledge” condition requires less information or effort and is thus less constrained. In practice the local knowledge information could be obtained from a pilot study such as a small scale presence-absence survey or from expert opinion.

The PBL method (Li et al., 2011) can predict the required probability well, but it is hard to gain the inferential relationship between the probability of a species’ presence and the influential environmental covariates. The CLK method, on the other hand, provides a remedy by finding the impact of the environmental covariates on species’ presence, whilst also providing good predictions. We also extend the local certainty or the prototypical presence location condition to the more general “local knowledge” condition. Similar to the PBL method, the performance of the refined CLK method is dependent on the classification method being chosen to fit the conditional labelling probability Pr(*s* = 1|*x*). In our numerical studies, a simple logistic regression as the classifier gave poor results when compared to the neural network (comparative results not shown). That is why we used the neural network in classifying Pr(*s* = 1|*x*) versus Pr(*s* = 0|*x*). Similar as other ML methods, our proposed approach also requires reasonably large background samples to enable reliable estimation, whereas the selection of large background samples can be easily achieved in practice.

An alternative suggestion by a referee is to perform/conceive the CLK method in a Bayesian framework, given its similarity to setting a highly informative prior (i.e. a spike prior) on the local knowledge at some sites. This could be a potential research to extend the CLK method in the framework of Bayesian analysis. Meanwhile it is worth mentioning that the conditional probability of ‘sampling/labelling’ *c* is assumed constant in our study. There has been research in both species distribution modelling and ML to study the case of non-constant *c*, such as modelling *c* as sampling bias through environmental covariates (Fithian et al., 2015). This suggests a potential direction for future research that would combine the ML and statistical methods to study non-constant c.

## 5 Conclusion

This paper firstly shows that the proper condition to estimate the absolute probability of presence for presence-only data is not the RSPF as the LK method has claimed. We found that the LK method is fragile and often fails to give reliable estimates even when the RSPF conditions are satisfied. Rather we propose the local knowledge condition, which relaxes the pre-determined population prevalence condition that has so often been used in much of the existing literature. The proposed concept of local knowledge extends the local certainty or the prototypical presence location assumption when the latter is not satisfied. The concept has significant implications for demonstrating the necessary condition to possibly estimate the absolute probability of presence without absence data in species distribution modelling.

## Acknowledgements

We thank the two anonymous referees for their very constructive feedback that has greatly improved the manuscript. This work was supported by Australian Research Council Grant No DP190100613.

## Authors’ Contributions

YW conceived the ideas and led the writings of the manuscript. CLS performed numerical analyses and led the writings of numerical study and Results sections. LS conceived the ideas and contributed to overall writings. All authors contributed critically to the drafts and gave final approval for publication.

## Data Availability

R scripts used to generate simulated data will be achieved at Github, should the manuscript be accepted.

## Appendix A: Relationship between the Absolute Probability of Presence and the Conditional Probability of Selection

The relationship is identified using Li et al. (2011) and Phillips et al. (2009). The definition of Pr(*s* = 1|*x*), i.e., the probability that a site will be chosen as a presence sample rather than a background sample, conditioned on the environmental variables (Phillips et al., 2009) is not similar to Pr(*y* = 1|*x*), i.e., the probability of occurrence conditioned on the environmental variables. We will now define the notations used.

We denote presence as *y* = 1 and absence as *y* = 0. Hence, the desired model can be written as P(*y* = 1|*x*). Let *π* = Pr(*y* = 1), the overall prevalence of the species of presence. and *p*_1_ is the number of observed presence data points in the presence sample in the training data set. The background data in the training set contain *p*_2_ presences and *n*_2_ absences, both in proportion to their overall population prevalence and *p*_2_ + *n*_2_ = *n*_0_. Moreover, let *s* = 1 denote the observed presence and *s* = 0 the background data. Remember that a background data point is either an unknown presence or an unknown absence. If *s* = 1, we know that *y* = 1; however, if *s* = 0, we do not know whether *y* = 1 or *y* = 0. Therefore,

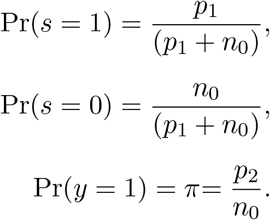

By definition of conditional probability, we have,

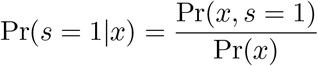

Then by applying Bayes rule, we have

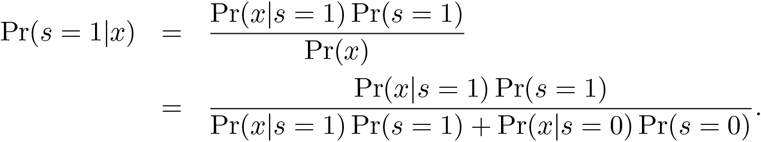

Then by definition of Pr(*s* = 1) and Pr(*s* = 0)

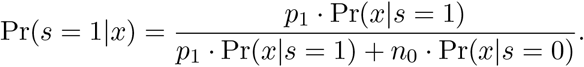

Next, dividing all terms by *p*_1_ · Pr(*x*|*s* = 1), we have

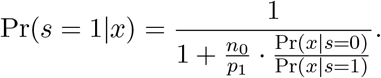

Pr(*x*|*s* = 1) is the density of the environmental covariates conditional on the observed presence, while Pr(*x*|*s* = 0) is the density of environmental covariates for the entire region. Therefore by definition, Pr(*x*|*s* = 0) = Pr(*x*) and Pr(*x*|*s* = 1) = Pr(*x*|*y* = 1).

By applying the Bayes rule again to Pr(*x*|*y* = 1) we get

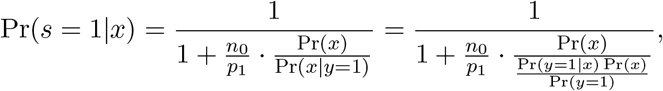

Hence,

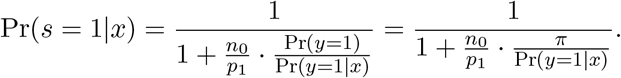

When the prototypical presence locations assumption is satisfied, i.e. Pr(*y* = 1|*x*) = 1, we can show that,

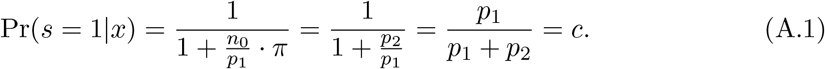

## Appendix B: Results of the LK method with larger samples size

**Figure B.1:**
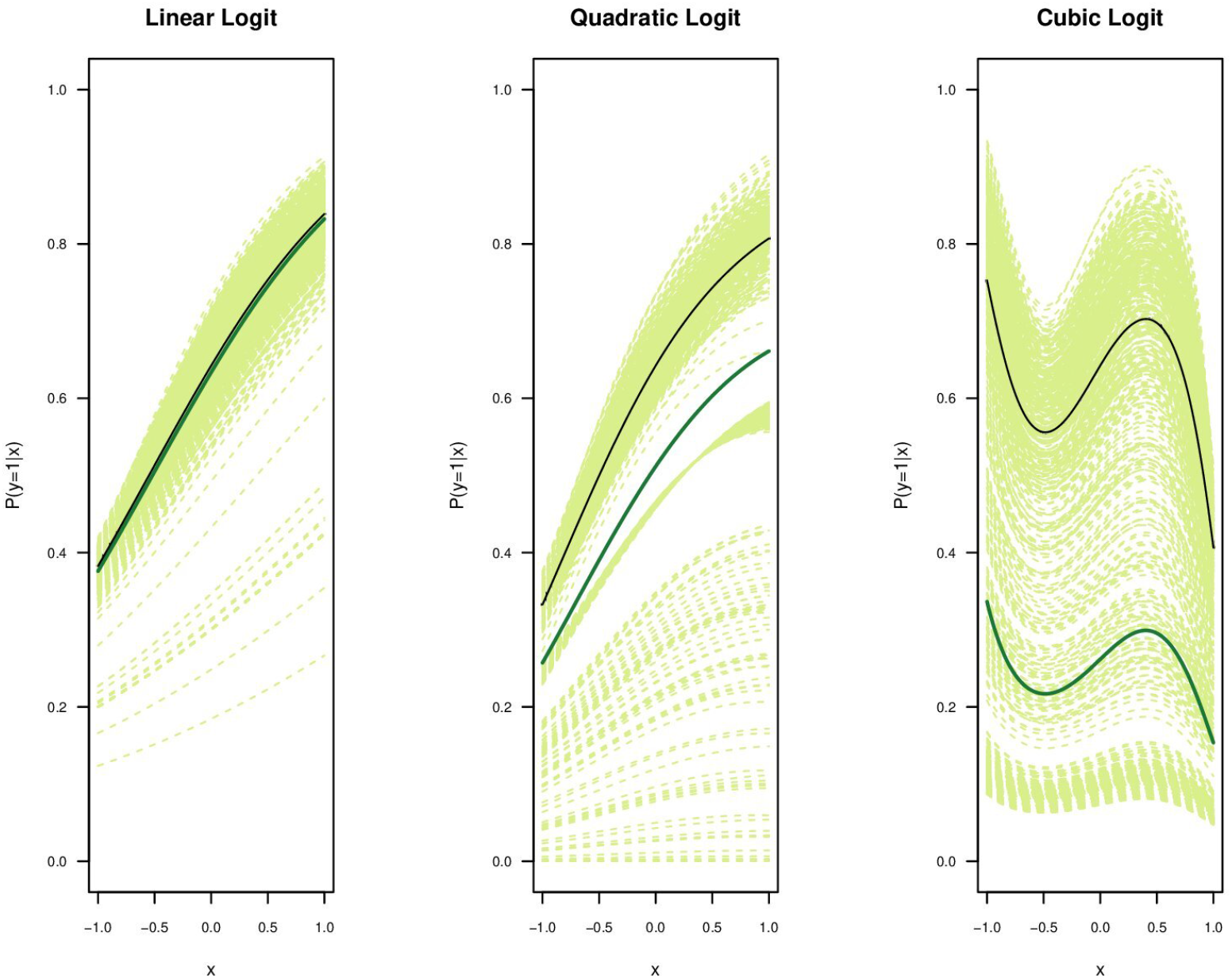
Category 3: The Pr(*y* = 1|*x*) is plotted when the original logistic parametric function only satisfies the RSPF condition (Category 3) for **very large sample size**. The original simulated function (black line), average LK method (green) is given with the respective 1000 simulation lines (light green) for the Linear Logistic (left), Quadratic Logistic (middle) and Cubic logistic (right). For each simulation, 50000 presence samples and 500000 background samples were chosen from a landscape with the predictor variable x in [−1, 1].

## Appendix C: Results of the complementary log-log (Cloglog) functions

The chosen cloglog functions are given in Figure C.1 and Table C1. These cloglog functions are divided into two categories, category one having three functions that satisfy both the RSPF and the local certainty condition and the second category only satisfies the RSPF conditions. These functions are also fitted using the logit parametric function to investigate the performance of the LK and CLK methods when the model is mis-specified. The results given in the Figs C.2 and C.3 show that the performance of the CLK method is good even though the models are mis-specified, whereas the LK method is still unable to accurately identify the original function as expected.

**Table C1:** Probability of presence for the simulated complementary log-log (Cloglog) species used in the experimental evaluation.

| Simulated Species | Probability of Presence |
| --- | --- |
| Category 1 | Both RSPF & Local Certainty Satisfied |
| Linear | $\Pr(y = 1\vert x) = 1 - \exp(-\exp(-0.886 - 1.16x))$ |
| Quadratic | $\Pr(y = 1\vert x) = 1 - \exp(-\exp(0.37 + 1.56x - 1.5x^2))$ |
| Cubic | $\Pr(y = 1\vert x) = 1 - \exp(-\exp(0.5855 + 1.064x - 0.218x^2 - 1.81x^3))$ |
| Category 2 | Only RSPF conditions Satisfied |
| Linear | $\Pr(y = 1\vert x) = 1 - \exp(-\exp(0.5855 + 1.064x))$ |
| Quadratic | $\Pr(y = 1\vert x) = 1 - \exp(-\exp(0.5855 + 1.064x - 0.218x^2))$ |
| Cubic | $\Pr(y = 1\vert x) = 1 - \exp(-\exp(0.1 - 0.064x - 0.85x^2 - 0.81x^3))$ |

**Figure C.1:**
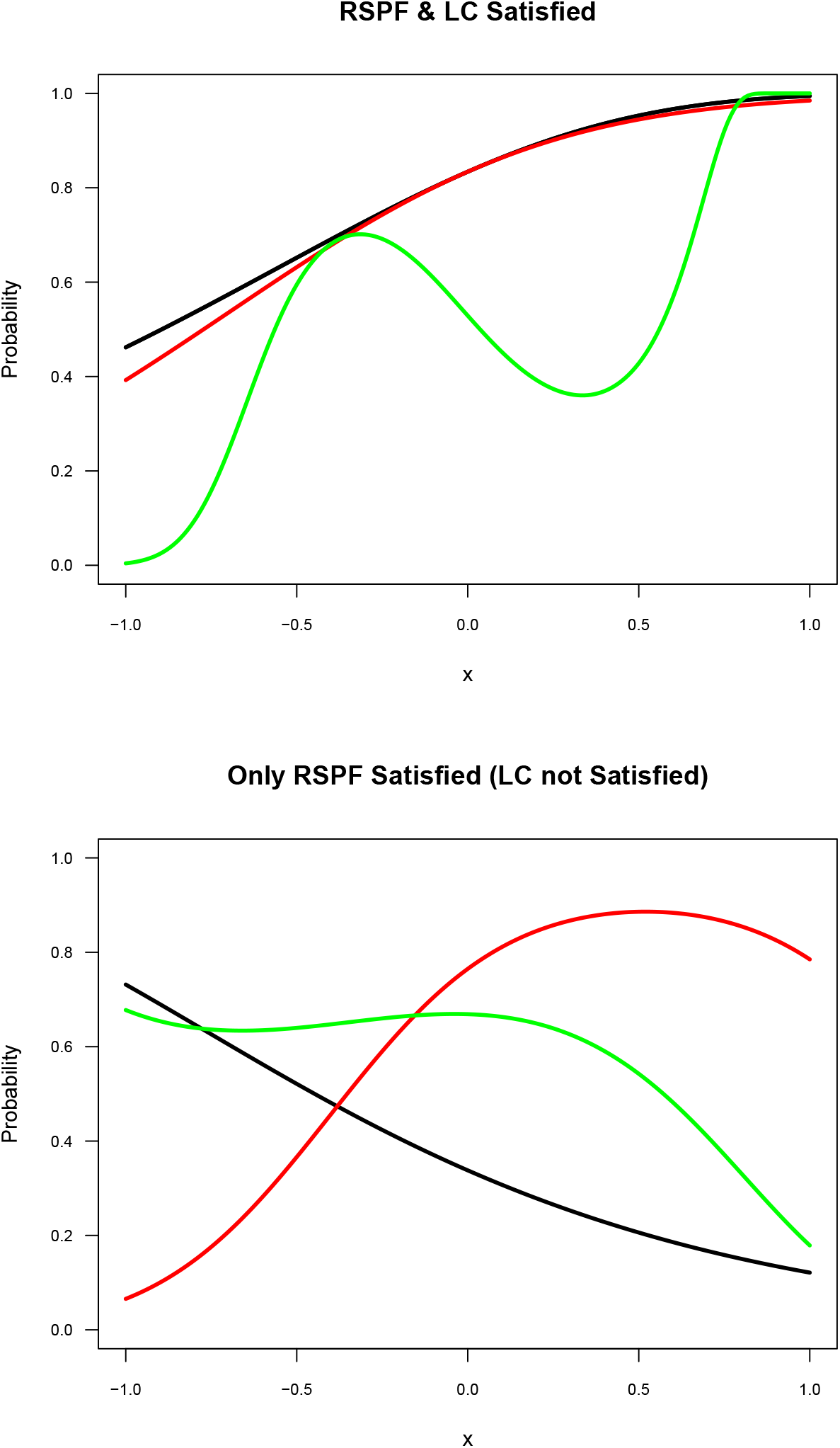
The selected simulated species distributions of the two groups of cloglog functions; category 1 and 2 respectively. The plots display the following functions named as follows. The top plot gives the functions of the category 1: only RSPF satisfied, that are linear (black), quadratic (red) and cubic (green). The bottom plot of category 2: RSPF and local certainty satisfied gives the linear (black), quadratic (red) and cubic (green).

**Figure C.2:**
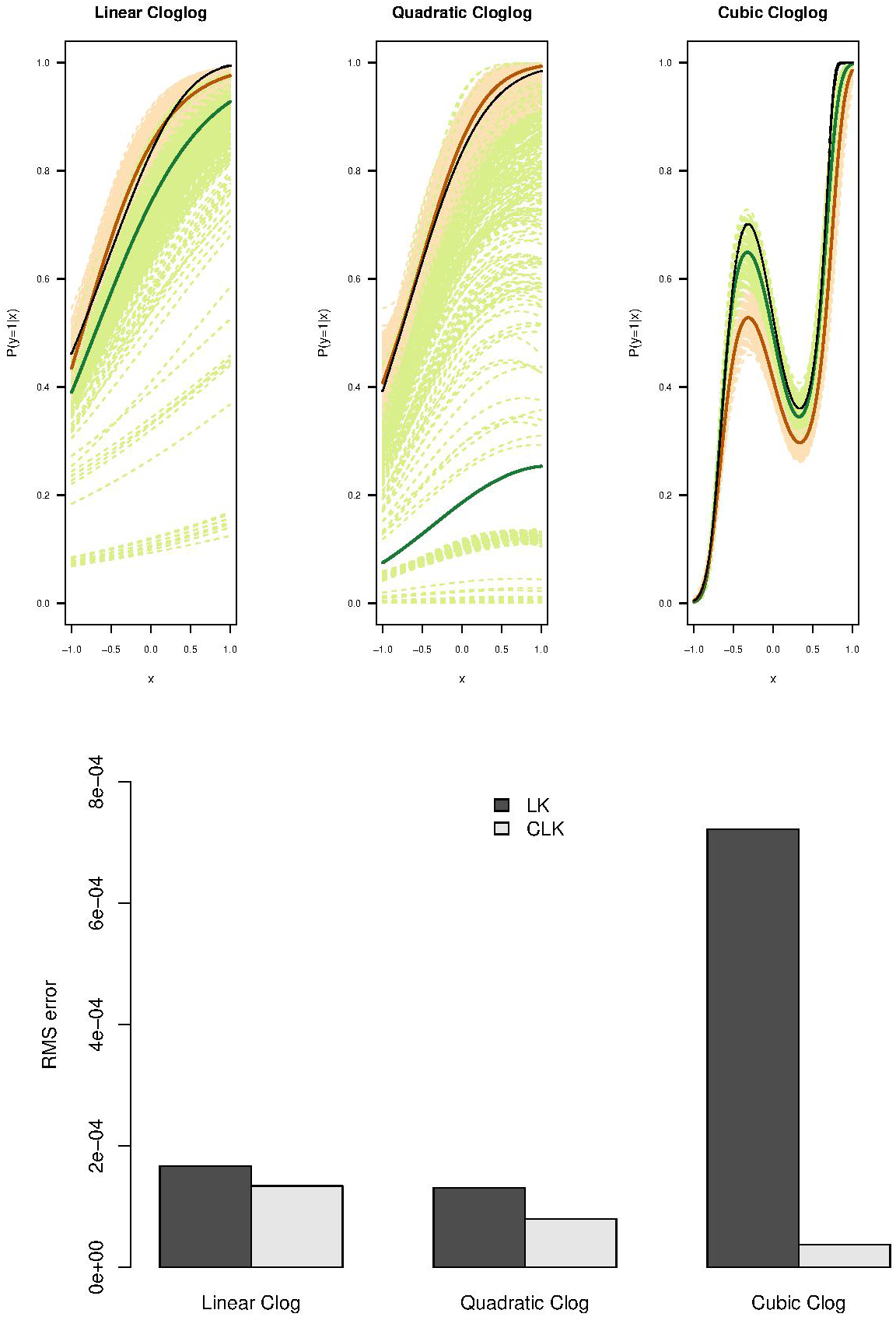
The Pr(*y* = 1|*x*) is plotted when the original Cloglog parametric functions that satisfy both the RSPF and the local certainty conditions as given by Solymos and Lele (2015). The original simulated function (black line), average LK method (green) and average CLK method (orange) are given with the respective 1000 simulation lines (light green dashed lines for LK and light orange dashed for CLK methods). For each simulation, 5000 presence samples and 50 000 background samples were chosen from a landscape with the predictor variable x in [−1, 1]. The bottom bar plot visualise the RMSEs of the two methods, LK and CLK for the above logistic functions.

**Figure C.3:**
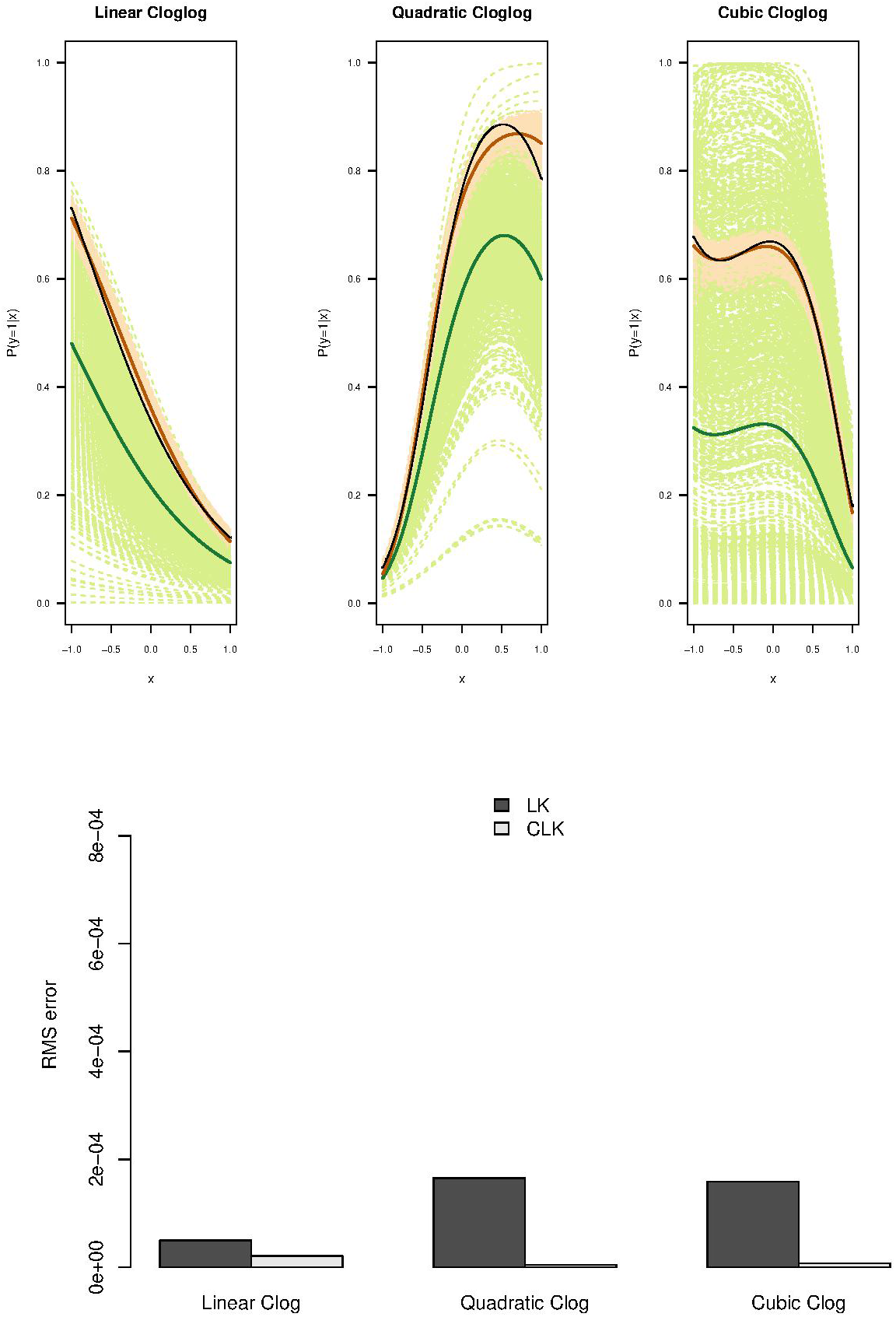
The Pr(*y* = 1|*x*) is plotted when the original Cloglog parametric functions that satisfy only RSPF conditions (with the local knowledge information assumed to be available as Pr(*y* = 1|*x*) = 0.73/0.89/0.68 for linear, quadratic and cubic functions respectively at maximum point of *x*). The original simulated function (black line), average LK method (green) and average CLK method (orange) are given with the respective 1000 simulation lines (light green dashed lines for LK and light orange dashed for CLK methods). For each simulation, 5000 presence samples and 50 000 background samples were chosen from a landscape with the predictor variable x in [−1, 1].The bottom bar plot visualise the RMSEs of the two methods, LK and CLK for the above logistic functions.

## Appendix D: Non-Convergence rate of LK method

These tables show the non-convergence rate of the LK method when fitting the models. It is also noted that the CLK method didn’t have any errors/non-convergences during model fitting.

**Table D. 1:**
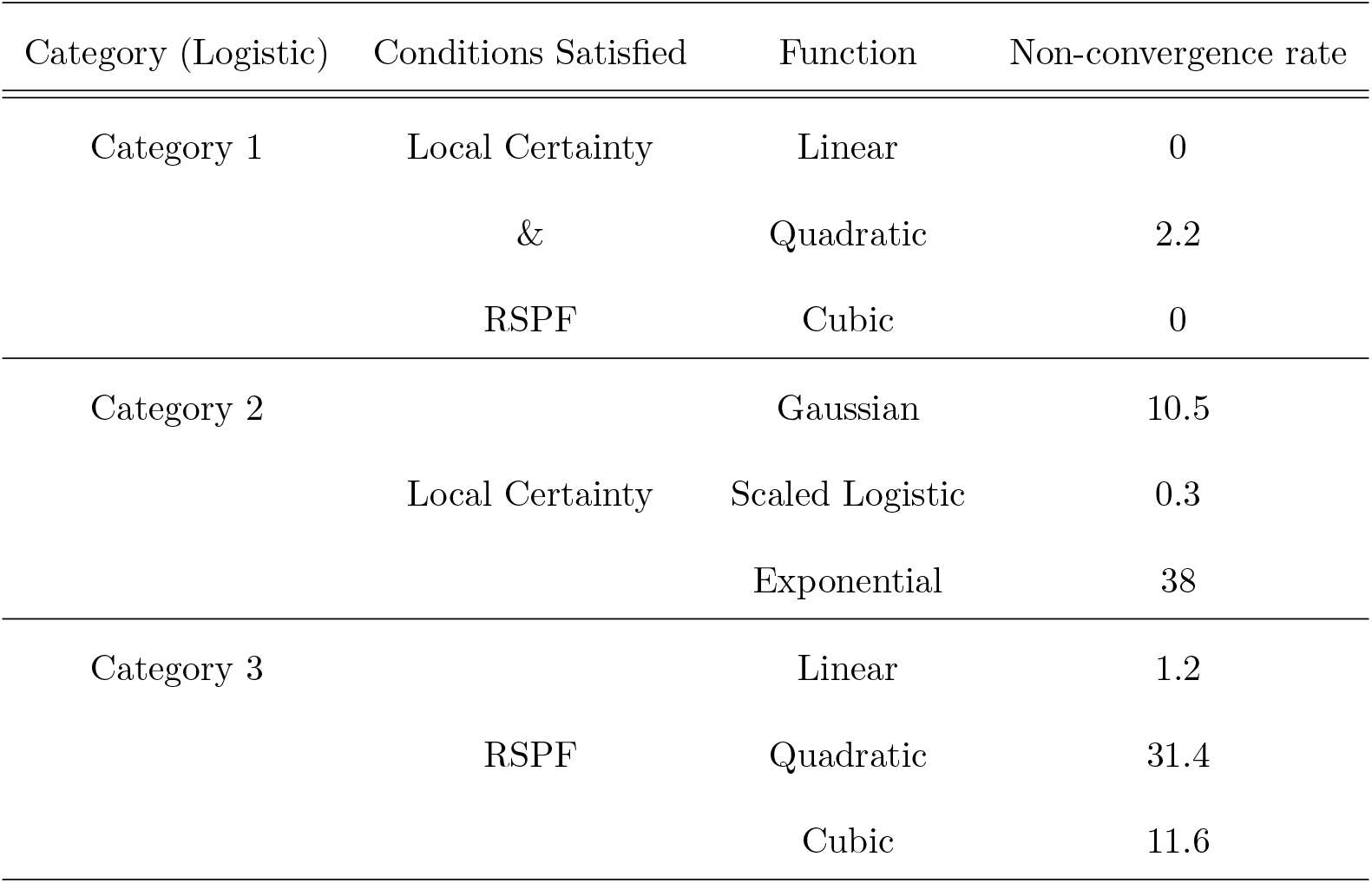
Proportion of non-convergence for the LK methods fitting three categories of logit functions with 1000 replications of simulations. 5000 presence samples and 50000 background samples were used in simulations.

**Table D. 2:** Proportion of non-convergence for the LK methods fitting two categories of complementary log-log (Cloglog) functions with 1000 replications. 5000 presence samples and 50 000 background samples were used in simulations.

| Category (Cloglog) | Conditions Satisfied | Function | Non-convergence rate |
| --- | --- | --- | --- |
| Category 1 | Local Certainty | Linear | 3.5 |
|  | & | Quadratic | 22.9 |
|  | RSPF | Cubic | 0 |
| Category 2 |  | Linear | 0.1 |
|  | RSPF | Quadratic | 1.5 |
|  |  | Cubic | 16.2 |

